# SANS ambages: phylogenomics with abundance-filter, multi-threading, and bootstrapping on amino-acid or genomic sequences

**DOI:** 10.1101/2025.03.20.643729

**Authors:** Fabian Kolesch, Marco Sohn, Andreas Rempel, Pia Hippel, Roland Wittler

**Affiliations:** Genome Informatics, Faculty of Technology and Center for Biotechnology, Bielefeld University, Bielefeld, 33615, Germany; Bielefeld Institute for Bioinformatics Infrastructure (BIBI), Bielefeld University, Bielefeld, 33615, Germany; Graduate School “Digital Infrastructure for the Life Sciences” (DILS), Bielefeld University, Bielefeld, 33615, Germany

**Keywords:** phylogeny, pangenomics, alignment-free

## Abstract

**Background:** The increasing amount of available genome sequence data enables large-scale comparative studies. A common task is the inference of phylogenies — a challenging task if close reference sequences are not available, genome sequences are incompletely assembled, or the high number of genomes precludes multiple sequence alignment in reasonable time. SANS is an alignment-free, whole-genome based approach for phylogeny estimation.

**Results:** Here we present a new implementation *SANS ambages* with a significantly increased application spectrum. It offers additional types of input data, parallelized processing, and bootstrapping. The source code (C++), documentation, and example data are freely available for download at: https://github.com/gi-bielefeld/sans. SANS can also be launched via the webinterface of the CloWM platform—free of charge, with a standard Life Science account: https://clowm.bi.denbi.de.

**Conclusions:** The new version not only shortens processing time on large datasets immensely by parallelization. Being able to also process amino acid sequences and offering a filter for low-abundant DNA read segments also enables new application cases. Bootstrapping and integrated visualization ease and enrich the interpretation of the resulting phylogenies.

## 1 Background

Comparative genomics often involves the reconstruction of phylogenies. The ever-increasing number of available genomes, many of which are published in an unfinished state or lack sufficient annotation, poses challenges to traditional phylogenetic inference methods that rely on the comparison of marker sequences. Whole-genome approaches still face the challenge that pairwise comparisons between all genomes result in a run time that increases quadratically with the number of input sequences– usually unsuitable in large-scale scenarios–whereas reference-based comparisons may introduce bias.

SANS [9, 11] is a whole-genome based, alignment- and reference-free approach that does not rely on a pairwise comparison of the genomes or their characteristics. In a pangenomic approach, evolutionary relationships are determined based on the similarity of the whole sequences. Sequence segments (*k*-mers) shared by a subset of genomes are interpreted as a *phylogenetic split* indicating the closeness of these genomes and their separation from the other genomes. The resulting splits can be visualized as a phylogenetic tree (Figure 1A) or network (Figure 1B). Both, the run time and the number of splits depends on the number of *k*-mers in the input. High sequence similarity among closely related genomes allows for a linear runtime in practice (Figure 2).

**Fig. 1.**
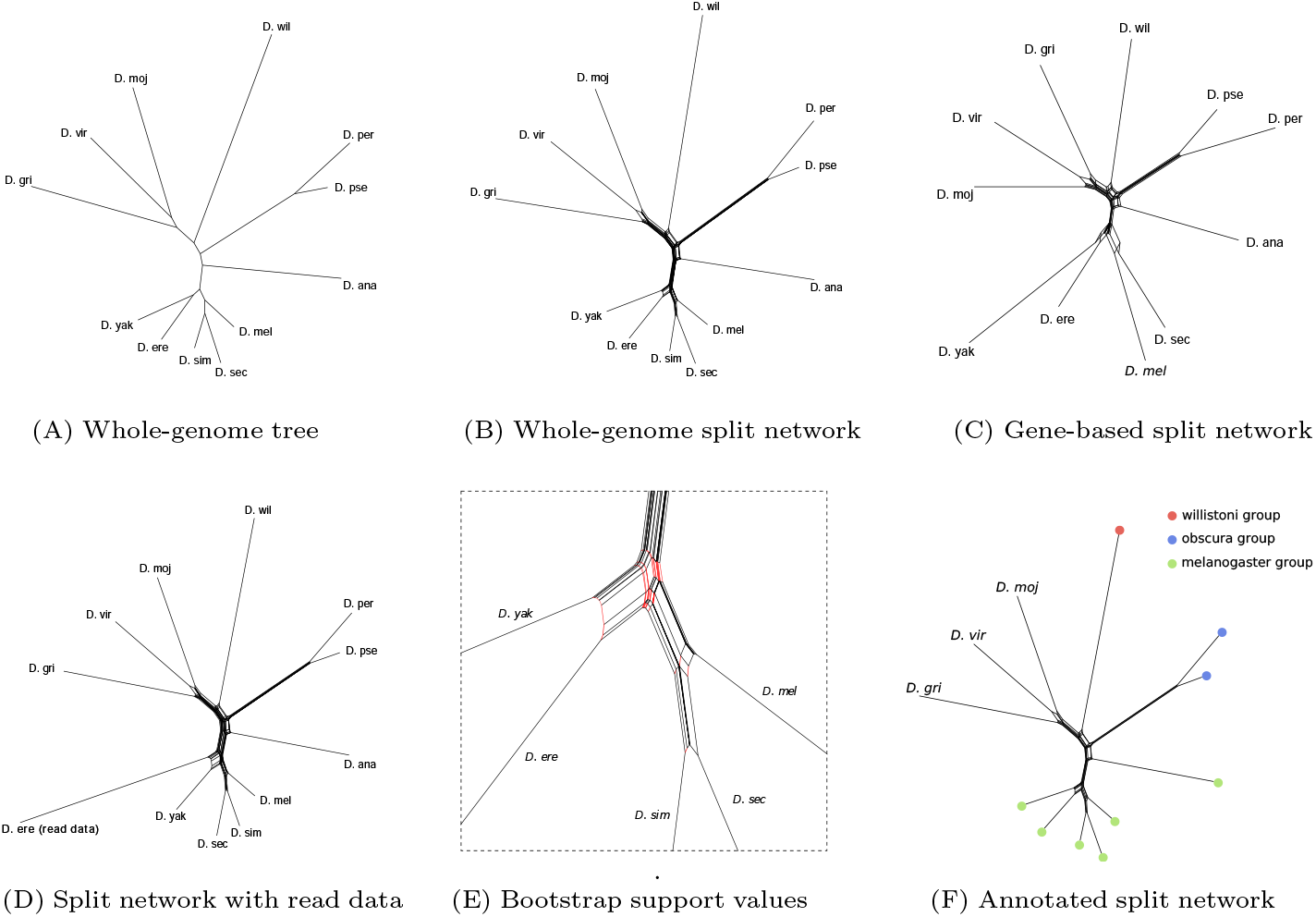
Demonstration of new features of SANS ambages. Phylogenies were visualized using Splits-Tree [2]. (A) Phylogenetic tree for twelve species of the genus *Drosophila*. SANS ambages was run on whole-genome data [1, files “all-chromosome” of release FB2023 03] with option `--filter tree` and default parameters otherwise. The tree topology agrees with the reference. (B) Phylogenetic split network on the same data as in Figure 1A obtained using `--filter weakly`. For possible interpretations of alternative splits, see [11]. (C) Phylogenetic split network on predicted gene sequences [1, files “all-predicted” of release FB2023 03]. For *D. simulans*, the corresponding file was not available. Translation and processing of amino acid sequences was called using `--code`. (D) Phylogenetic split network on the same data as in Figure 1A, but with one genome, *D. erecta*, replaced by long read data [8, SRR7167960]. Option --qualify 3 was used to filter for *k*-mers appearing at least three times in the read data, whereas a minimum of 1 was set for the assembled data. (E) Section from Figure 1B in which red edges indicate a support value below 75 % based on 1000 replicates (`--bootstrap 1000`). (F) Phylogeny from Figure 1B with the taxa of subgenus *Sophophora* assigned to their respective group using option `--label`.

**Fig. 2.**
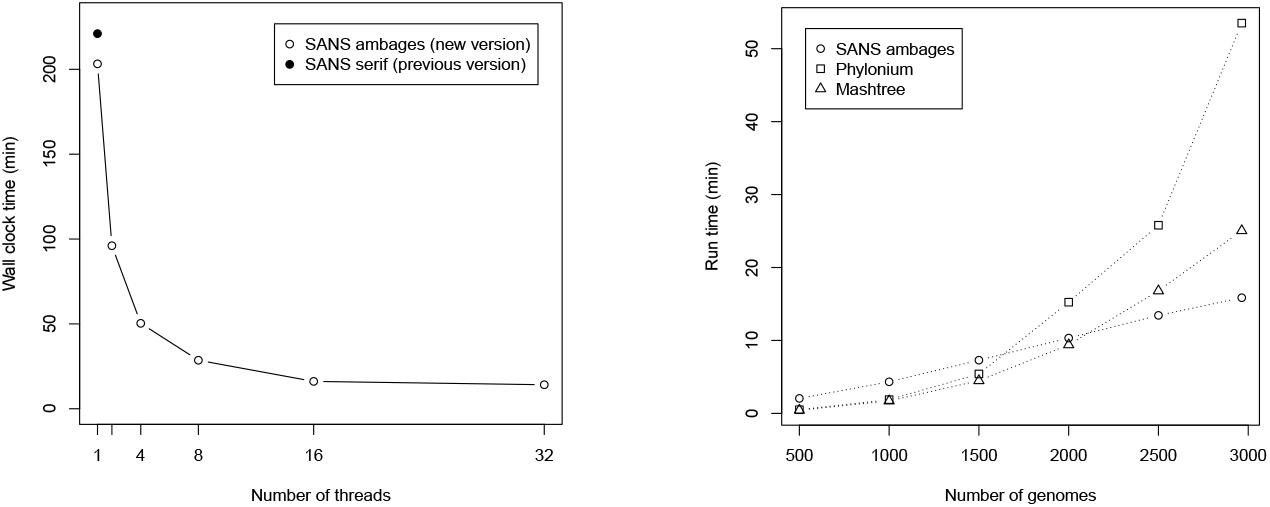
Run times (wall clock) of SANS ambages measured on 2964 *Salmonella enterica* subspecies *enterica* assemblies [9, 12] on a ∼ 2 GHz virtual machine with 32 cores and 1 TB of RAM. (Left) The run time for varying numbers of threads was evaluated using option `--threads` and also compared to the previous version SANS serif without multithreading support (git commit 3053ffb5). (Right) Run time comparison to Phylonium [5, version 1.7] with mattools (https://github.com/evolbioinf/mattools, commit 3af4fc8), and Mashtree [4, version 1.4.6] with Mash [7, version 2.2]. All tools were run on 16 threads with default parameters; SANS ambages was run with filter `-f strict` to obtain a tree.

## 2 Implementation

Here, we present the new version SANS ambages (**a**bundance-filter, **m**ulti-threading, and **b**ootstrapping on **a**mino-acid or **ge**nomic **s**equences) with several substantial improvements. Besides processing DNA sequences (whole genomes or assemblies), SANS ambages can also work on amino acid level, taking protein sequences (translated or untranslated) as input. Further, the ability to process read data has been enhanced by the option to filter out low-abundant sequence segments. Bootstrapping allows to augment the output with confidence values or to compile phylogenies with high-confidence splits. The integration of SplitsTree [2] and additional visualization options allow automated graphical output. Finally, SANS ambages is considerably more performant than its predecessor [9]. In particular, multiple input sequences can now be processed in parallel, which will be elaborated and evaluated in the following.

SANS ambages supports (gzipped) Fasta and Fastq files as input and is capable of indexing the *k*-mers of multiple input files in parallel. Each thread pops a file from a list of unread files and moves a window of length *k* over the sequences. Similar to the previous version of SANS, extracted *k*-mers are indexed as a key, with a bit vector to represent the set of input files in which the *k*-mer is present as a value. To avoid issues due to concurrent access to the bit vectors, the space of *k*-mers is split into *m* partitions, each represented by a sub-index. Each sub-index can be accessed by one thread at a time, ensuring thread safety and exact results. To keep the probability of two threads trying to access the same sub-index small, the number of partitions is chosen magnitudes higher than the number of threads. To compute the partition a *k*-mer belongs to, its binary representation is interpreted as an unsigned integer modulo *m*. Similar to a rolling hash function, the partition can be updated in constant time when shifting from one *k*-mer to the next.

The above indexing is preceded by two filtering steps. Firstly, if an abundance threshold is given (see Section 3.2), each *k*-mer is temporarily stored in an additional data structure before it is further processed. For a threshold of 2, a set data structure is used to store *k*-mers that are seen exactly once so far in the processed genome. For larger thresholds, a counter is maintained for each *k*-mer until it passes the abundance filter. Secondly, a considerable amount of *k*-mers is unique to individual genomes and storing their occurrence explicitly takes less memory compared to their occurrence bit vectors. Thus, each *k*-mer is first stored in a hash map, together with the identifier of the genome it has been observed in. If it is observed in a second genome, it is transferred to the final *k*-mer occurrence table.

The run time depends on both the total number of *k*-mers in the input and the number of splits. On the one hand, the maximum number of splits is exponential in the number of input genomes. But on the other hand, it is also bounded by the number of *k*-mers, because each *k*-mer can induce at most one split. In practice, for closely related sequences, high redundancy allows for a linear run time trend (Figure 2).

## 3 Results

In the following, we elaborate on the new abilities of SANS and demonstrate their use. For the sake of visualization, we selected a rather small dataset of up to twelve species of the genus *Drosophila* for which different kinds of input data (whole-genome, gene predictions, long-reads) as well as a commonly accepted reference phylogeny were available [1, 8]. Some improvements are illustrated on a larger dataset of about 3000 *Salmonella enterica* subspecies *enterica* assemblies [9, 12], for which a thoroughly determined phylogeny is available as a reference [12, Figure 2A, supertree 3].

### 3.1 Amino acid sequences as input

The new version enables the processing of protein sequences, either translated, or untranslated employing automatic translation including different genetic codes. By switching from whole-genome to protein sequences, intergenic regions are omitted or the input is even restricted to core genes. Furthermore, the length of the amino acid sequences is only one-third of the corresponding nucleotide sequences. On the one hand, this reduction of input sequence implies some loss of phylogenetic signal. On the other hand, the larger alphabet size, tuning out silent mutations, and higher conservation in coding regions yield a clearer signal. We observed that using amino acid sequences exhibits similar accuracy compared to using DNA sequences. For instance, for the *Salmonella* dataset, running SANS ambages on gene predictions (obtained by Prodigal [3] v2.6.3) yields higher accuracy with respect to the reference phylogeny [12, Figure 2A, supertree 3] than using the whole-genome data (F1 score under Robinson-Foulds metric of about 91 % and 82 %, respectively).

A side effect of the reduced amount of input that needs to be processed, is a substantial reduction of the computational resources. For instance, for the *Salmonella* dataset (using parameters `--filter weakly` and 16 threads), run time was reduced by 69 % (to 13 minutes) and the memory peak by 80 % (to 9 GB). For the *Drosophila* dataset, run time was reduced by 63 % (to 4 minutes) and the memory peak by 49 % (to 16 GB). The resulting phylogeny is shown in Figure 1C.

### 3.2 Abundance filter for read data

When analyzing read data, a common preprocessing step is to filter out low coverage *k*-mers that typically arise from sequencing errors. SANS ambages includes a new option to perform such a filtering step, allowing raw read data to be analyzed without the need to run another tool first. A minimum coverage threshold can be specified, i.e., `--qualify` *n* filters out all *k*-mers that occur less than *n* times per genome. With the new filtering option, SANS ambages is now able to process large read datasets that were impossible to analyze with the previous version in a reasonable amount of memory. For example, the unfiltered analysis of a subset of ten genomes in a *Drosophila* long-read dataset [8] could not be completed on a system with 1 TB of RAM, while the same analysis using the filter option `--qualify` 2 required only 119 GB of RAM. It is also possible to define the minimum coverage value individually for each genome in the list of input files, making it possible to use both reads and assembled genomes in the same analysis pipeline. For this purpose, SANS ambages is now able to parse the *file of file* format, which was previously used by the tool *kmtricks* [6]:

~~~
<Identifier> : <File1> ; … ; <FileN> ! <MinCoverage>
Taxon1 : /path/to/assembled_genome.fasta ! 1
Taxon2 : ./reads_fw.fastq.gz ; ./reads_rv.fastq.gz ! 3
~~~

This is particularly useful when adding a newly sequenced genome, i.e., read data, to an existing reference dataset, allowing it to be placed in a known phylogeny without the need for assembly or other processing. This scenario is mimicked in the phylogeny shown in Figure 1D.

### 3.3 Bootstrapping

To assess the robustness of phylogenetic signals with respect to noise in the input data, bootstrap replicates are constructed by randomly varying the observed *k*-mer content. If the overall input contains *n* distinct *k*-mers, we simulate the process of drawing *n k*-mers uniformly at random with replacement. Then, a phylogenetic split that is originally supported by *x k*-mers is supported by *x* times *p k*-mers, where *p* is sampled following the binomial distribution describing the expected number of times an individual *k*-mer is drawn. Each such bootstrap replicate is then further processed like the original dataset, and for each split its relative abundance among the replicates is determined as bootstrap support. There is also the option to construct a consensus split set based on the support values, which is then annotated with the split weights of the original split set. Bootstrap replicates are constructed and processed in parallel. Figure 1E shows a section of the phylogenetic split network in which red edges indicate bootstrap support values below 75 %. These splits of low confidence in fact disagree with the reference tree [1].

### 3.4 Visualization

Visualizing the phylogeny provides an intuitive overview and facilitates interpreting the determined splits. SANS ambages therefore offers the Nexus file format as output, which can be opened in SplitsTree (version 4) [2] for an interactive visualization. A PDF file can also be automatically created via executing SplitsTree in the background.

The new option `--label` allows to label a phylogeny. Taxa can be assigned to groups by providing a tab-separated file:

~~~
<Identifier> <Group>
Taxon1 Group_A
Taxon2 Group_B
Taxon3 Group_A
~~~

In the visualization, the taxon IDs will then be replaced by colored circles according to their respective group. The colors are either selected automatically or can be provided by the user in an additional tab-separated file. Figure 1F shows an example of a partially labeled phylogeny.

### 3.5 Run time improvement

Run time improvements are illustrated on the *Salmonella* dataset of about 3000 assemblies [9, 12]. See Table A1 for further exemplary datasets.

As can be seen in Figure 2 (Left), in comparison to the previous version, run time on a single thread is reduced by 8 % (from 221 to 203 minutes). Peak memory usage is reduced by 34 % (from 62 GB to 41 GB, data not shown). Using two threads, run time is reduced further by 52 %, using 0.4 % more memory. Increasing the number of threads from one to 32, run time is reduced by 93 % to 14 minutes, using 14 % more memory (46 GB). The observation that doubling the number of threads yields a run time reduction of more than 50 % can be explained by a commonly known effect in parallelization: more compute cores also possess more cache and thus improve memory access [10].

Figure 2 (Right) shows a linear run time trend for an increasing number of input sequences. Compared to Mashtree [4] and Phylonium [5], SANS has the highest run time of 2 minutes on 500 sequences but is fastest on the complete data set. The reconstruction accuracy of all tools with respect to a reference phylogeny [12, Figure 2A, supertree 3] are comparable as shown in Table A3.

## 4 Conclusions

We presented SANS ambages for alignment- and reference-free phylogenetic inference. This new version offers substantial improvements. New types of input data can be analyzed: On the one hand, a filter for low-abundant DNA sequences enables to use read data instead of or in combination with assembled sequences. On the other hand, protein sequences, such as raw gene predictions or selected core genes, can be processed. Bootstrap support values and simplified visualization facilitate the interpretation of the phylogenetic split networks. Effective parallelization makes it possible to process a multiple of input files in the same time.

## Availability and requirements

*Project name:* SANS

*Project home page:* https://github.com/gi-bielefeld/sans *Operating system(s):*

Platform independent *Programming language:* C++ version 17

*Other requirements:* To visualize the splits, we recommend the tool SplitsTree [2].

*License:* SANS ambages is licensed under the GNU general public license. Integrated libraries are licensed under the MIT, BSD-2, and LGPL license.

*Any restrictions to use by non-academics:* None.

## Declarations

### Ethics approval and consent to participate

Not applicable.

### Consent for publication

Not applicable.

### Availability of data and materials

Sources for the datasets analysed during the current study are listed in Appendix A, Table A2.

### Competing interests

The authors declare that they have no competing interests.

### Funding

This project has received funding from the European Union’s Horizon 2020 research and innovation programme under the Marie Sklodowska-Curie agreement [grant number 872539] and the BMBF-funded German network for bioinformatics infrastructure (de.NBI) [grant 031A533].

### Authors’ contributions

FK developed and implemented the parallelization, MS implemented the amino acid functionality, AR implemented the abundance filter, PH implemented the visualization options, RW developed and implemented the bootstrapping. All authors read and approved the final manuscript.

## Acknowledgements

The authors acknowledge the Bielefeld-Gießen Center for Microbial Bioinformatics (BiGi) of the BMBF-funded German network for bioinformatics infrastructure (de.NBI) for provision of compute resources and general support.

## Appendix A Overview of exemplary datasets

**Table A1.**
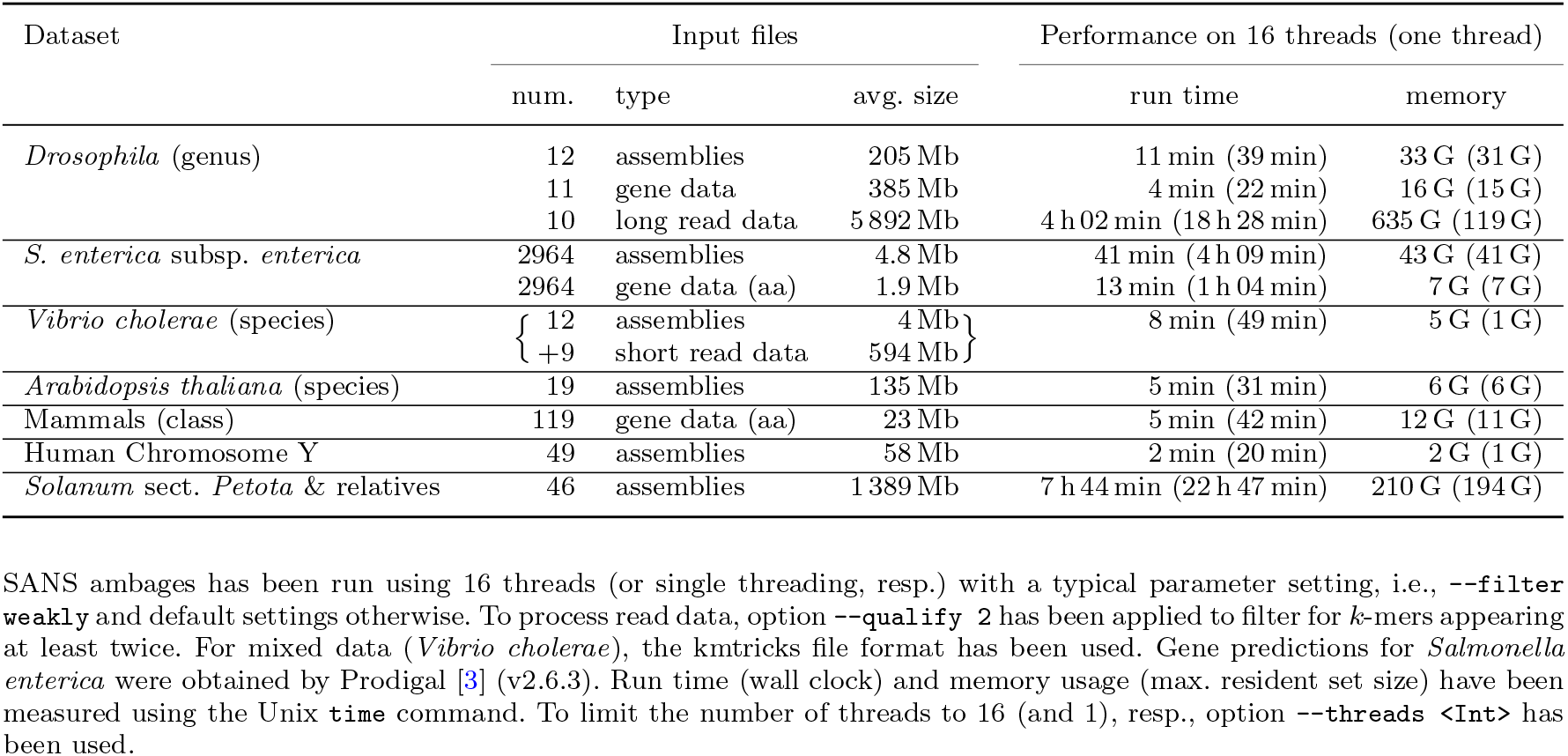
Overview of exemplary datasets and the required computational resources.

**Table A2.**
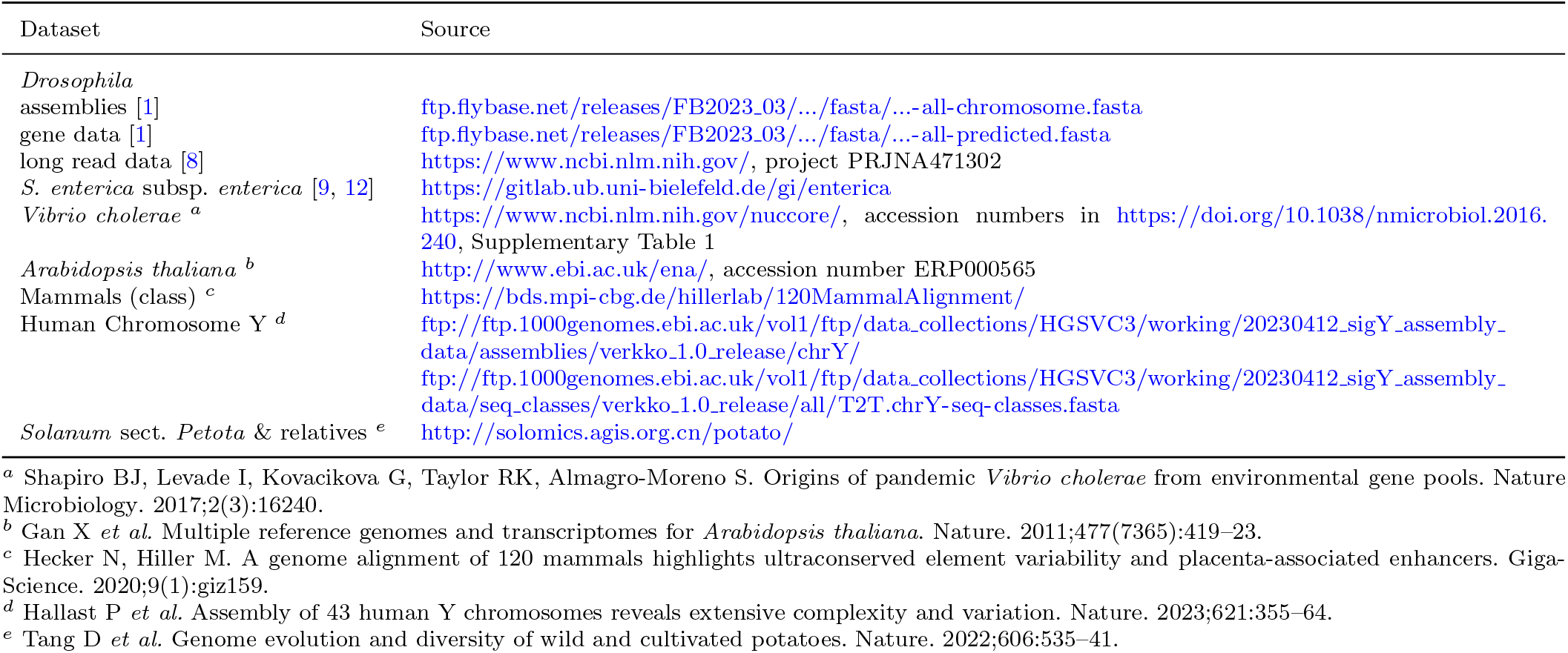
Links to datasets listed in Table A1.

**Table A3.** F1 score (harmonic mean of precision and recall) of splits called by the different tools on the *S. enterica* subsp. *enterica* dataset, as well as with respect to the reference phylogeny [12, Figure 2A, supertree 3]. All tools have been run with default parameters, SANS ambages was run with filter -f strict to obtain a tree. Neighbor Joining was used to compute a tree from the distances obtained by Phylonium.

|  | Phylonium | Mashtree | Reference tree |
| --- | --- | --- | --- |
| SANS ambages | 0.72 | 0.72 | 0.79 |
| Phylonium |  | 0.71 | 0.75 |
| Mashtree |  |  | 0.72 |

## Notes

### Competing Interest Statement

The authors have declared no competing interest.

